# Identification of new bacterial type III secreted effectors with a recursive Hidden Markov Model profile-alignment strategy

**DOI:** 10.1101/081265

**Authors:** Xi Cheng, Xinjie Hui, Aaron P. White, Zhirong Guo, Yueming Hu, Yejun Wang

## Abstract

To identify new bacterial type III secreted effectors is computationally a big challenge. At least a dozen machine learning algorithms have been developed, but so far have only achieved limited success. Sequence similarity appears important for biologists but is frequently neglected by algorithm developers for effector prediction, although large success was achieved in the field with this strategy a decade ago. In this study, we propose a recursive sequence alignment strategy with Hidden Markov Models, to comprehensively find homologs of known YopJ/P full-length proteins, effector domains and N-terminal signal sequences. Using this method, we identified 155 different YopJ/P-family effectors and 59 proteins with YopJ/P N-terminal signal sequences from 27 genera and more than 70 species. Among these genera, we also identified one type III secretion system (T3SS) from *Uliginosibacterium* and two T3SSs from *Rhizobacter* for the first time. Higher conservation of effector domains, N-terminal fusion of signal sequences to other effectors, and the exchange of N-terminal signal sequences between different effector proteins were frequently observed for YopJ/P-family proteins. This made it feasible to identify new effectors based on separate similarity screening for the N-terminal signal peptides and the effector domains of known effectors. This method can also be applied to search for homologues of other known T3SS effectors.

## Introduction

Bacterial type III secretion systems (T3SSs) play important roles in interaction with host cells. ^1, 2^ T3SSs are widely distributed among gram-negative bacteria; more than one hundred genera have been found with at least one copy of T3SS (Hu et al., unpublished). The apparatus component proteins of a T3SS assemble a needle-like conduit spanning bacterial cellular membranes, recognizing and translocating T3SS effector proteins into host cells. ^1, 2^ These effector substrates exert the pivotal function to facilitate the interaction between bacteria and host cells, further causing symbiosis or pathogenesis. Disclosing molecular interactions between the effectors and host cellular molecules as well as their consequences, would facilitate understanding corresponding bacteria-host interaction mechanisms.

It is consistently a big challenge to identify new type III secreted effectors. ^3, 4^ Effectors are frequently transferred among bacterial strains by horizontal transfer and the number and catalog of effectors varies between species. Moreover, the nucleotide and amino acid sequences show low similarity and few conserved features among different effectors, making it difficult to identify new ones by sequence alignment strategies. Despite some debate, overwhelming experimental and bioinformatic evidence supports the hypothesis that the N-terminal amino acid sequences of effectors contain T3SS-recognized signals. ^5–9^ This opens a window for in silico screening for new effectors by bioinformatic methods. Various conserved properties exist in the N-termini of effectors, and based on these features, dozens of algorithms have been developed to make de novo predictions. ^8–14^

Success was achieved using the established bioinformatic algorithms and tools, but with large limitations. High false positive rates are the main problem, which is caused by over-fitting of the models. Currently, methods tend to be too focused on the sequence and structural property of N-terminal signals of effectors, while other regions (e.g., effector domains, chaperone binding domains, etc.) and other important features (e.g., evolutionary property of species or sequences) were seldom strengthened or combined for consideration. ^15–18^ Some hypotheses also need to be further justified, for example, the common signal recognition mechanisms among T3SSs and the minimum signal length. These hypotheses formed the basis for the various effector prediction algorithms. After justification, the training effectors should be further refined and categorized, conserved features are trained more specifically and precisely, and the predictors are finally built with an expectation of higher accuracy. We have been in such an attempt to build a T3SS effector prediction package. According to our and others’ experience, the sequence alignment-based strategies, if possible, always generated more precise results. ^19–21^ These effector candidates should be screened with a higher priority in the first stage. However, it is often difficult to find proteins with low homology to known effectors. In this research, we focused on a T3SS effector family, YopJ/P, exploring its evolutionary sequence properties. YopJ/P-family effectors are distributed in multiple human or plant pathogens, playing important roles in bacterial interaction with host cells as a ubiquitin-like protein protease. ^22–24^ We expected to use YopJ/P as an example, showing how to better use the global and regional sequence conservation and Hidden Markov Models to find with a higher sensitivity more probable new effectors in multiple bacterial strains.

## Results

### 1 Variation of the signal sequence and conservation of the effector domain

Through global sequence alignment with blastp and a full-length protein HMM, a number of YopJ/P homologs were disclosed from *Yersinia*, *Aeromonas*, *Vibrio*, *Edwardsiella*, *Escherichia*, *Lonsdalea*, *Serratia*, *Bartonella*, *Salmonella*, *Candidatus Hamiltonella*, *Arsenophonus* and *Pseudovibrio* (Table 1; Supplemental File 1). Among these genera, *Yersinia*, *Aeromonas*, *Vibrio*, *Edwardsiella*, *Escherichia*, *Lonsdalea* and *Salmonella* were reported with at least one copy of T3SS. Our previous research screened T3SSs from *Lonsdalea*, *Serratia*, *Candidatus Hamiltonella*, *Arsenophonus* and *Pseudovibrio*, respectively (Hu et al., unpublished). For *Bartonella*, the homologs are of high similarity with *Yersinia* YopJ/P, and are present in a variety of species and strains (Table 1). However, Hidden Markov Models (HMMs) based on ten core apparatus proteins did not detect any T3SS from these strains.

**Table 1:** YopJ/P homologs based on global alignment of full-length proteins1.

| Genus | Genome | Species \| Subspecies | Note |
| --- | --- | --- | --- |
| *Yersinia* | Plasmid | *Y. enterocolitica* | Gene Symbol: yopP \| yopJ |
|  |  | *Y. pestis* |  |
|  |  | *Y. pseudotuberculosis* |  |
|  |  | *Y. aldovae* |  |
|  |  | *Y. ruckeri* |  |
| *Aeromonas* | Plasmid | *A. salmonicida* subsp. *achromogenes* | Gene Symbol: aopP |
|  |  | *A. salmonicida* subsp. *salmonicida* |  |
|  |  | *A. salmonicida* subsp. *masoucida* |  |
| *Vibrio* | Chromosome | *V. sagamiensis* | Gene Symbol: vopP |
|  |  | *V. cholerae* |  |
|  |  | *V. parahaemolyticus* |  |
| *Edwardsiella* | Unclear | *E. piscicida* | Only in one piscicida strain |
|  |  | *E. tarda* | In 3 *E. tarda* strains |
| *Escherichia* | Unclear | *E. coli* | In 3 *E. coli* strains |
| *Lonsdalea* | Unclear | *L. quercina* | Only in an unknown strain |
| *Serratia* | Unclear | *S. symbiotica* | Only in an unknown strain |
| *Bartonella* | Chromosome | *B. taylorii*, *B. washoensis*, | No T3SS detected |
|  |  | *B. florencae*, *B. rattaustraliani*, |  |
|  |  | *B. alsatica*, *B. vinsonii*, |  |
|  |  | *B. tribocorum*, *B. grahamii*, |  |
|  |  | *B. elizabethae*, *B. quintana*, |  |
|  |  | *B. rattimassiliensis*, *B. doshiae*, |  |
|  |  | *B. clarridgeiae*, *B. rochalimae* |  |
| *Salmonella* | Chromosome | *S. bongori* | Gene Symbol: avrA |
|  |  | *S. enterica* subsp. *enterica* |  |
|  |  | *S. enterica* subsp. *diarizonae* |  |
| *Hamiltonella* | Chromosome | *Candidatus Hamiltonella defensa* | Only in an unknown strain |
| *Arsenophonus* | Unclear | *A. nasoniae* | Only in an unknown strain |
| *Pseudovibrio* | Chromosome | Unclassified | In two strains |

Note: The protein accessions and sequences were listed in Supplementary File 1.

In most of the genera, YopJ/P homologs are present in a limited subset of species or strains. For *Yersinia*, *Salmonella* and *Bartonella*, multiple species/strains encode the gene (Table 1). In *Yersinia*, *yopJ/P* is encoded on plasmids, and mobile elements like integrases are often found to flank the gene (Figure 1). The *avrA* gene, encoding the YopJ/P homolog in *Salmonella*, is located in a pathogenic island (Figure 1). The *yopJ* gene in *Bartonella* is flanked by phage proteins and therefore is likely originated from a phage (Figure 1). Taking together, the *yopJ/P* genes are widely spread, apparently through various horizontal gene transfer events.

**Figure 1.**
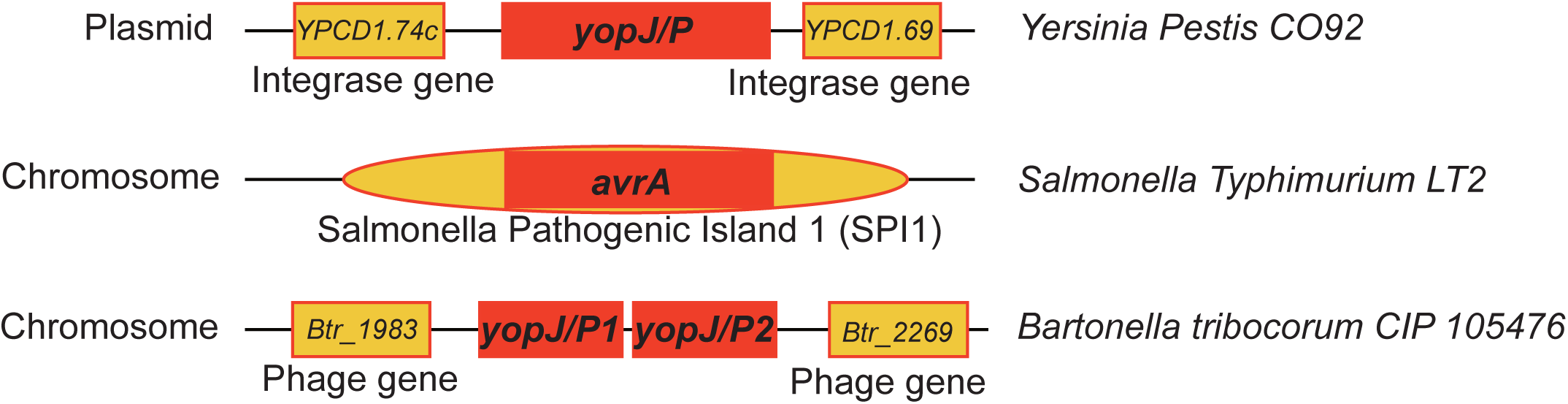
Horizontal gene transferring signature of yopJ/P genes. The yopJ/P gene was shown in red box in each genus; the genome was shown in black line; the adjacent genes or pathogenic island region were shown in golden blocks.

The similarity between the full-length YopJ/P homologs of different bacteria is uneven for different regions of the proteins (Figure 2). Generally, the N-terminal signal sequences show large variations, and the C-terminal effector domains are conserved. For each pair of YopJ/P homologs, the N-terminal 60-aa peptide fragments are most divergent, while the C-terminal effector domains are highly conserved, and the N-terminal 61~120-aa peptide fragments show a medium similarity level between the flanking regions but closer to the C-terminal core effector domains (Figure 2).

**Figure 2.**
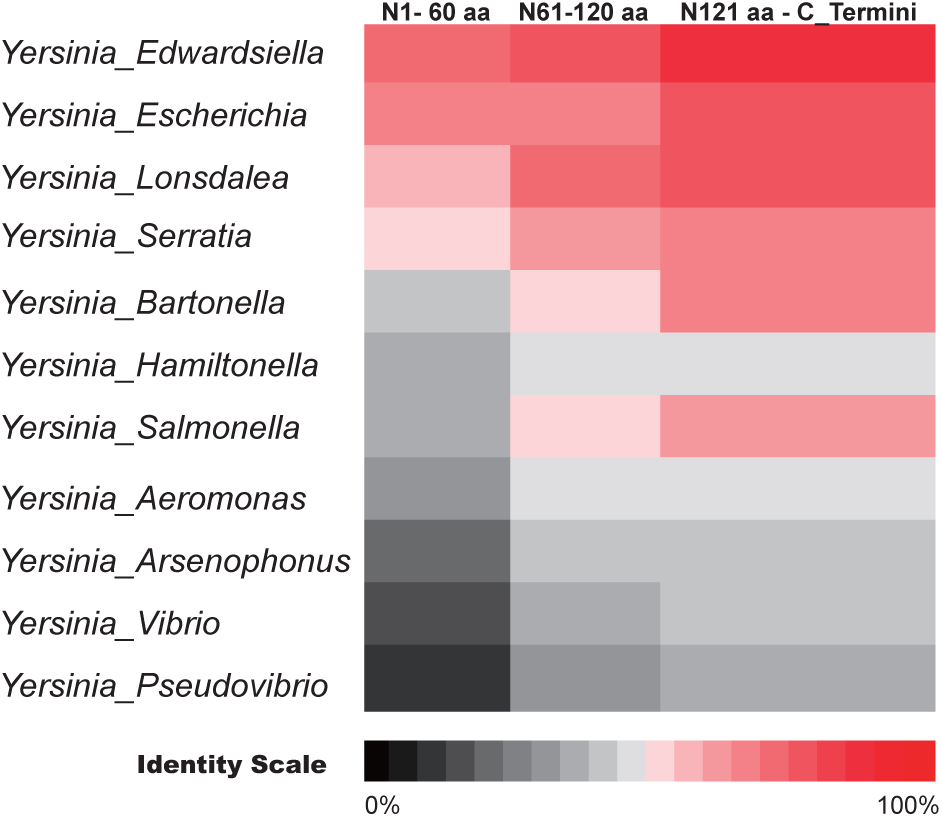
The divergence of N-termini and conservation of effector domains. The average YopJ/P sequence similarities (identities) between Yersinia and other genera were shown in different color. The sequences were compared for three sub-regions: N-terminal 1-60 amino acids, 61-120 amino acids, and 121-C terminus.

### 2 Effectors identified by sequence similarity of C-terminal effector domains

Based on previous reports and the current observations on the distribution of pairwise similarities along sub-fragments of YopJ/P proteins, the N-terminal ~100-aa region was defined as signal peptide while the rest part represented the YopJ/P effector domain. The effector domains were extracted and used for HMM profile building for two steps, which were furthermore used for new YopJ/P effector screening (Materials and Methods; Supplemental File 1).

Besides the homologs identified by global alignment of full-length YopJ/P proteins, new effectors were screened from *Pseudomonas*, *Burkholderia*, *Candidatus Regiella*, *Uliginosibacterium*, *Acidovorax*, *Xanthomonas*, *Ralstonia*, *Rhizobacter*, *Xenorhabdus*, *Marinomonas* and *Sinorhizobium* (Table 2). These proteins also frequently show evidence of horizontal transfer signatures, with flanking transposition enzyme encoding genes or phage genes. Except *Uliginosibacterium*, *Rhizobacter* and *Xenorhabdus*, the other genera were identified with at least one copy of T3SS in representative strains. Core apparatus genes based HMMs identified one T3SS from *Uliginosibacterium* and two T3SSs from *Rhizobacter* but none from *Xenorhabdus* (Figure 3).

**Figure 3.**
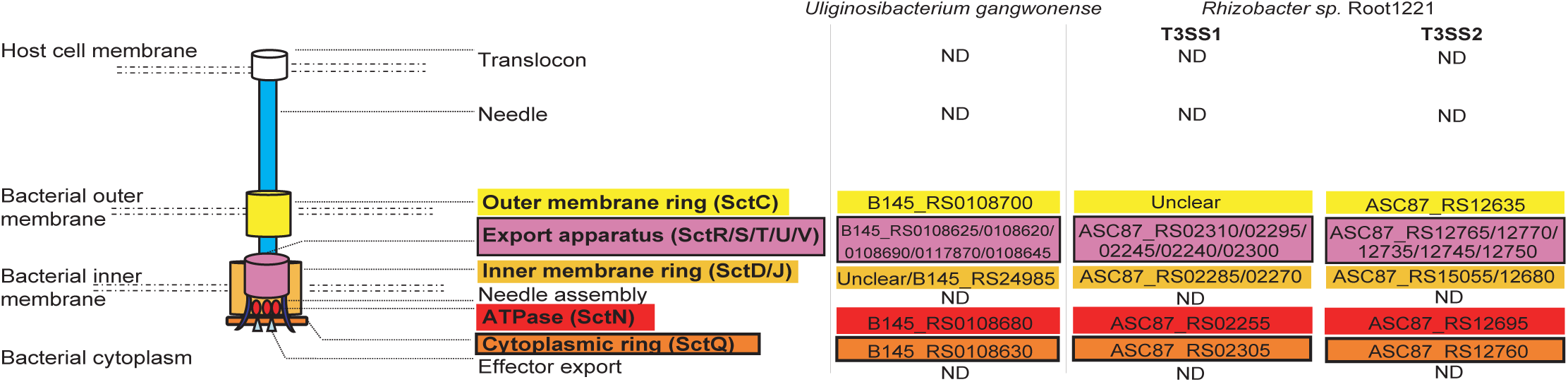
New T3SSs identified from *Uliginosibacterium* and *Rhizobacter*. Each apparatus component and its encoding proteins were shown in different color. For the T3SSs of *Uliginosibacterium gangwonense* and *Rhizobacter sp. Root1221*, the accession for each apparatus component encoding protein was shown. ‘Unclear’ accessions represented the genes not detectible from the current genome; ‘ND’ meant ‘not detected’.

**Table 2:** Proteins conserved for the YopJ/P effector domain but varied for the N-terminal sequences1.

| Genus | Genome | Species \| Subspecies | Note |
| --- | --- | --- | --- |
| *Pseudomonas* | Chromosome/<br>Plasmid | *P. syringae*, *P. avellanae*<br>*P. amygdali*, *P. savastanoi*,<br>*P. tremae*, *P. cannabina*,<br>*P. coronafaciens* | Gene Symbol: hopZ4 \| avrPpiG1 \|<br>hopZ2 \| hopZ1 \| orf34 \| hopPmaD \|<br>hopZ3 |
| *Regiella* | Chromosome | *Candidatus Regiella*<br>*insecticola* | Only in a strain 5.15 |
| *Uliginosibacterium* | Chromosome | *U. gangwonense* | Only in a strain DSM 18521 newly<br>detected with a T3SS |
| *Acidovorax* | Chromosome | *A. citrulli*, *A. avenae*<br>*A. oryzae* |  |
| *Xanthomonas* | Chromosome | *X. campestris*<br>*X. euvesicatoria*<br>*X. gardneri*<br>*X. translucens*<br>*X. axonopodis* | Gene Symbol: xopJ \| xopJ2 \|<br>avrRxv \| avrBsT |
| *Rhizobacter* | Chromosome | *Unclassified* | Only in strain *R. sp.* Root1221<br>newly detected with 2 T3SSs |
| *Burkholderia* | Chromosome | *B. dolosa*<br>*B. andropogonis* | Only in a dolosa strain AU0158<br>and an andropogonis strain<br>ICMP2807 |
| *Marinomonas* | Chromosome | *M. mediterranea* | Only in a strain MMB-1 |
| *Sinorhizobium* | Plasmid | *S. fredii* | Gene Symbol: nopJ \| y4IO |
| *Ralstonia* | Chromosome | *R. solanacearum*<br>*R. pickettii* | Gene Symbol: avrRxv |
| *Xenorhabdus* | Chromosome | *X. bovienii*<br>*X. cabanillasii*<br>*X. khoisanae*<br>*X. doucetiae* | In multiple strains not detected<br>with T3SS |

Note: The protein accessions and sequences were listed in Supplementary File 1.

In *Pseudomonas*, the YopJ/P-family effectors are only present in species that represent plant pathogens (Table 2). There are multiple homologs with the YopJ/P effector domain, including AvrPpiG1, Orf34, HopPmaD, HopZ1, HopZ2, HopZ3 and HopZ4 encoded by multiple species (Table 2). The protein sequences vary widely in the length and composition of the N-terminal signal region; however, many of the proteins have been verified to secrete through a T3SS conduit. Two *Burkholderia* species, human pathogen *B. dolosa* and plant pathogen *B. andropogonis*, either has a strain showing a YopJ/P effector domain containing protein (AJY11376.1 and KKB61318.1) respectively (Table 2). The two proteins vary a lot between each other for the full-length proteins, and both showed a relatively conserved YopJ/P effector domain and strikingly varied N-terminal peptide sequences. Insect symbiont *Candidatus Regiella* 5.15, nematode symbiotic or pathogenic *Xenorhabdus* strains (*X*. *bovienii* Intermedium, *X. cabanillasii* JM26, *X. khoisanae* MCB and *X. doucetiae* FRM16), an environmental *Uliginosibacterium gangwonense* strain DSM 18521, strains of three plant pathogenic *Acidovorax* species (*A. citrulli*, *A. avenae* and *A. oryzae*) and multiple plant pathogenic *Xanthomonas* and *Ralstonia* species, plant symbionts *Rhizobacter sp.* Root1221 and *Sinorhizobium fredii* NGR234, and a marine bacterium *Marinomonas mediterranea* MMB-1, each also has one or more N-terminal divergent YopJ/P-containing proteins (Table 2). Some of the homologs in *Xanthomonas*, *Ralstonia* and *Sinorhizobium* have been proved to be true effectors delivered to host cytoplasm through T3SS conduits.

### 3 Splitting and fusion of N-terminal signal sequences

In some bacteria, especially *Pseudomonas*, *Xanthomonas* and other plant pathogenic species, the YopJ/P proteins show an interesting N-terminal splitting and fusion phenomenon, which was previously noted as ‘terminal reassortment’. ^17^ CAJ23833.1 and AJW66323.1 encode two putative T3SS effectors XopJ and AvrRxv1, respectively, both of which are YopJ/Pfamily members. ^25,26^ However, CAJ23833.1 and AJW66323.1 showed strikingly different N-terminal signal sequences (Figure 4A). Compared with EKQ58453.1, which also encodes a putative T3SS effector protein XopJ1, CAJ23833.1 shows high homology but has apparent N-terminal extension. Similarly, CAJ22102.1 and KFA04636.1 show good alignment against AJW66323.1 but both have apparent N-terminal curtailment or extension (Figure 4A). CAJ22102.1 and KFA04636.1 also encode putative T3SS effectors AvrRxv2 and AvrRxv3, respectively.

**Figure 4.**
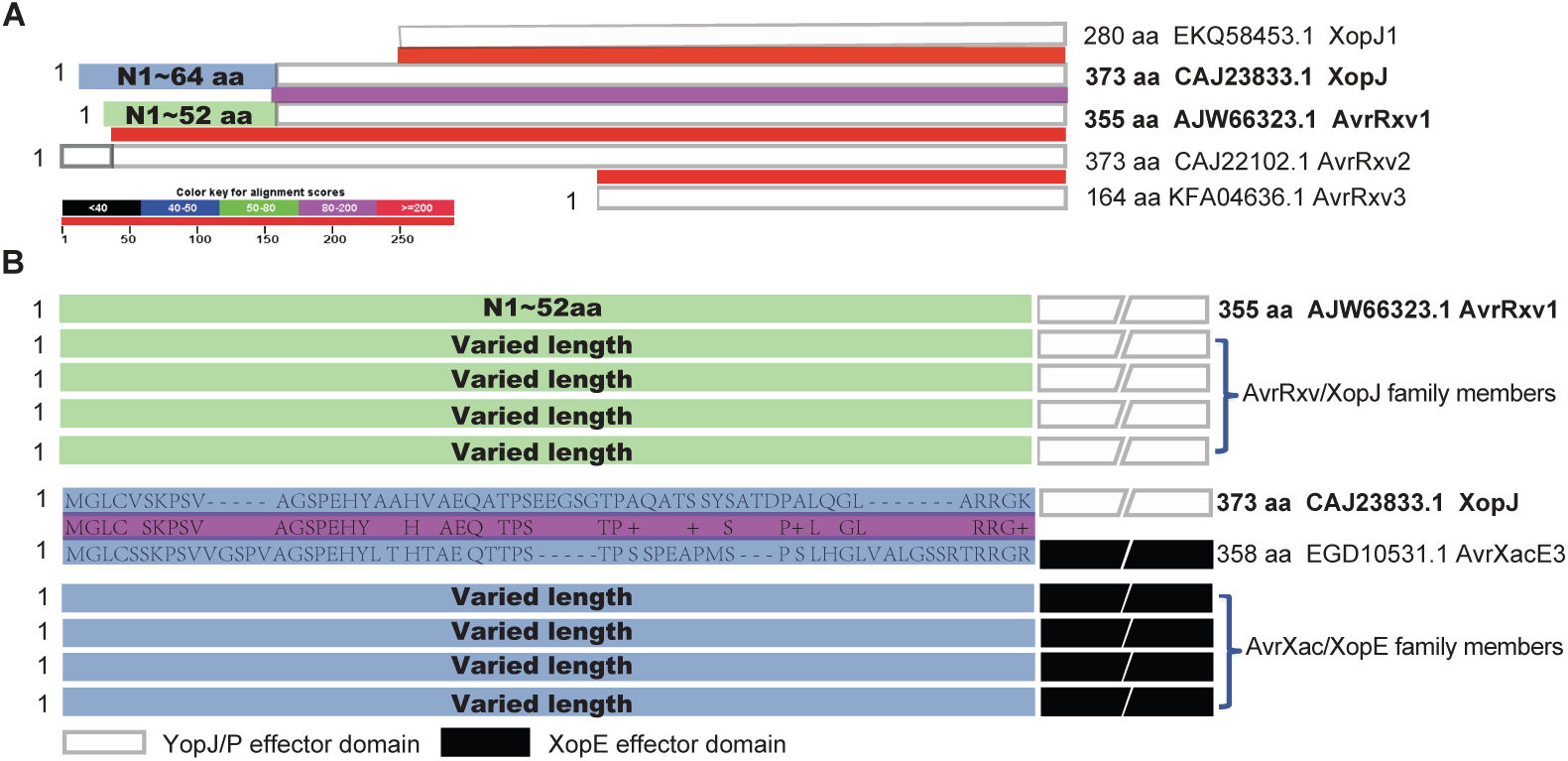
Extension, splitting and re-fusion of N-terminal signal sequences of T3SS effectors. (A) The exchange of N-terminal signal sequences of XopJ and AvrRxv1 and the N-terminal curtailment or extension of XopJ or AvrRxv protein family. Each protein was represented as a rectangular box and the left terminal colored boxes of the YopJ and AvrRxv1 proteins represented corresponding N-terminal signal sequences. The colored box between each vertically adjacent pair of proteins indicated the sequence similarity. The similarity score scale was shown in the left lower corner. (B) The diagram showing the proteins or protein families with the homologous N-terminal signal sequence of AvrRxv1 or XopJ. The left-end colored box represented the signal regions while the broken white or black box represented the YopJ/P or XopE effector domains. The purple box between XopJ and AvrXacE3 showed the similarity level and the alignment between the signal sequences of the two proteins. The N-termini of AvrRxv1 and XopJ were shown in the rectangular box in the same color for (A), respectively.

Homologs of the N-terminal signal sequence of AJW66323.1 were found to widely fuse with divergent YopJ/P effector domains (Figure 4B). For the N-terminal signal sequences of CAJ23833.1, homologs were not only found to fuse with YopJ/P effector domains, but also fused to other domains, such as XopE effector domains (Figure 4B). These AvrXacE/XopE family proteins were validated to be true effectors secreted through *Xanthomonas* T3SS conduits. ^26^ Similarly, the signal sequences of N-terminal extended AJW66323.1 or N-terminal reduced CAJ23833.1 homologs were also found to fuse with different YopJ/P or XopE effector domains.

Taken together, the results indicated that the T3SS signal sequences could be considered as domain-like units, which may fuse with or split from other domains as a whole to enable or disable the translocation of effectors via T3SS conduits.

### 4 Identification of new T3SS effectors with YopJ/P N-terminal homology profiles

The homologs of different N-terminal signal sequences of YopJ/P proteins were collected to build HMMs, which were used in a two-step recursive procedure to identify potential new effectors with T3SS-recognizable signals (Materials and Methods; Supplemental File 2).

In total, 4 major profiles and 8 minor profiles were classified for the N-termini of YopJ/P proteins. The largest profile comprised protein fragments from *Yersinia*, *Salmonella*, *Aeromonas*, *Edwardsiella*, *Serratia*, *Vibrio*, *Pseudovibrio*, *Escherichia* and *Lonsdalea*. The sequences showed a significant number of substitutions but insertions or deletions were rare. In contrast, for *Bartonella, Yersinia* and others, despite high similarity for the full-length proteins or effector domains, the N-termini of YopJ/P often showed large or multiple insertions and deletions. Because no T3SS was detected in strains with genome sequences finished (as shown before), the *Bartonella* YopJ/P N-termini were excluded for profile training. The other 3 major profiles were mainly present in plant pathogens or symbionts.

Though most proteins had been identified through global sequence alignment or YopJ/P effector domain profile screening, multiple effectors from 7 different genera were newly identified using the N-terminal signal profiles (Table 3). Four genera, including *Rhizobium*, *Mesorhizobium*, *Pantoea* and *Erwinia*, were newly identified with YopJ/P homologs or YopJ/P signal-profile effectors.

**Table 3:**
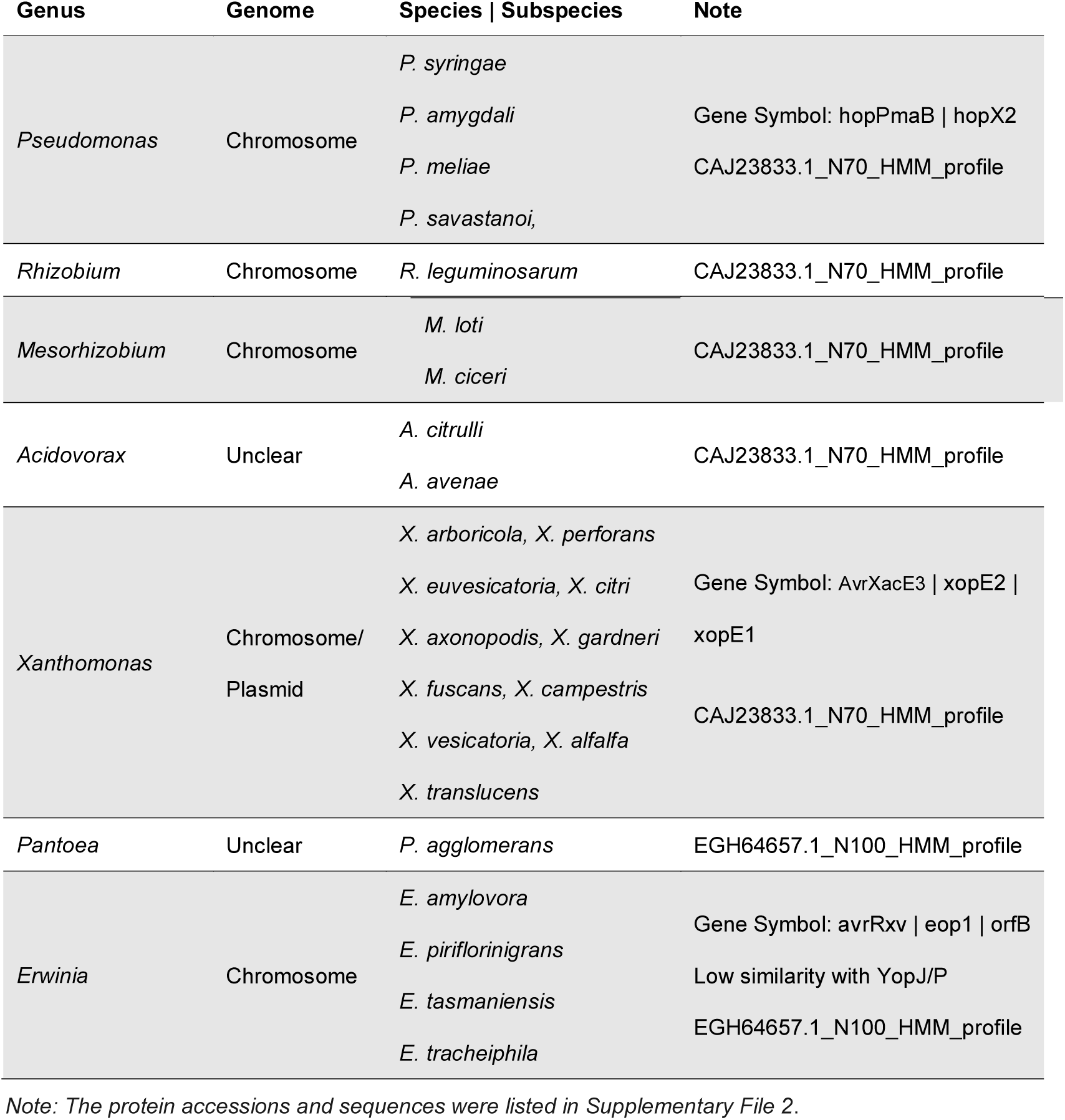
New effector candidates identified by sequence homology for N-terminal signal sequences of YopJ/P proteins.

Taking together, we used Hidden Markov Models to screen the homologs of YopJ/P full-length proteins, effector domain and N-terminal signal sequences in a recursive way, and found hundreds of new and non-identical T3SS effector proteins from 27 genera and more than 70 species (Supplementary Files 1 and 2).

## Discussion

In this research, we proposed a recursive Hidden Markov Model profile-alignment strategy to identify new T3SS effectors based on homology to the validated YopJ/P proteins, C-terminal effector domains and N-terminal signal regions. Compared with other weakly conserved features at hierarchical levels, sequence similarity appeared more direct, frequently indicating functional conservation and giving highly confident prediction results. Different feature-learning algorithms and machine learning models becoming popular in T3SS effector prediction recently stems from an opinion that T3SS effectors are too divergent in sequence and traditional methods based on sequence alignment could hardly identify new effectors. In fact, however, more real new effectors have been identified by cloning experiments rather than by machine learning algorithms. It is noted that earlier bioinformatic methods achieved greater success in identification of T3SS or other effectors. These methods are no more than sequence alignment based. Therefore, an ideal effector screening strategy could be to find out all the homologs of known effectors (with high confidence to be true effectors), followed by machine learning algorithms for looking for possible full-new candidates (with tentative new clues but also with high false positive rates). In practice of the first step, the blast-based sequence alignment method seemed difficult to discover the distant homologs. In this report, we took YopJ/P family as an example to show a new recursive HMM-based sequence alignment strategy, finding a batch of new T3SS effectors. The method could be easily applied to other effector family. The HMM profiles built here are also useful for effective effector screening in new genomes.

Sequence similarity generally predicts the conservation of function. Therefore, the effectors identified through homology screening strategies are expected to be true with higher probability. However, we found putative YopJ/P members in *Bartonella* and *Xenorhabdus*, two genera for which representative strains were not found with any T3SS apparatus. *Bartonella* is particularly interesting. For a majority of *Bartonella* species, each has at least one copy of YopJ/P homolog, and a selection analysis demonstrated a strong purifying selection of the gene (data not shown). Since there was no T3SS identified in *Bartonella*, it’s unlikely that the YopJ/P protein could exert its function as a T3SS substrate. It is possible that the protein might have other routes to enter host cells to exert effector function, or perhaps it has other important roles inside bacterial cells. It is noted that the N-termini of *Bartonella* YopJ/P proteins showed apparent inserts and deletions (indels), whereas the homologs in other genera like *Salmonella*, *Aeromonas*, *Edwardsiella*, *Escherichia*, *Yersinia* and others, seldom had indels even though the global sequence similarity was not as high. Even though a very limited subset of strains had YopJ/P proteins in many genera other than *Bartonella* and *Xenorhabdus*, these strains all had at least a T3SS, and therefore, the proteins still had the ability to translocate through the T3SS conduit and function as T3SS effectors. The genes could be acquired through horizontal transfer events since there were often signatures shown in the flanking sequences (Figure 1).

We found three T3SSs in two bacterial genera, one in *Uliginosibacterium* and two in *Rhizobacter*. ^27–28^ *Uliginosibacterium gangwonense* DSM 18521, isolated from a wetland, was not clear if it could invade any host. Since a T3SS and at least one effector were identified, it is most likely that this bacterium has the capability to invade and interact with host cells. ^27^ *Rhizobacter sp*. Root1221 is a member of *Arabidopsis* root microbiota and therefore could probably interact with *Arabidopsis* root cells through the two copies of T3SSs. ^28^ Although some bacteria such as *Bartonella* and *Xenorhabdus* obtained some effectors but do not have any T3SS, a strain detected with a putative effector is more likely to have T3SSs. Therefore, the de novo identification of effectors could also indicate the higher presence probability and prompt the identification of T3SSs in the strain.

In this research, we took the YopJ/P-family proteins as an example, to show that the effector domain itself could be used for effector screening, and that the splitting and re-fusion of T3SS signal sequences and different effector domains help identify more effectors by looking for the N-terminal homologs. Effectors of different families from the same or different bacterial species were frequently found with low sequence similarity, some showed structural or functional homology, and they were called ‘T3-orthologs’ together. ^29^ We showed here a relatively larger variance in the N-terminal signal sequences without which more effector domains could show higher and detectible similarity (Figure 2). On the other hand, Stavrinides et al showed the ‘terminal reassortment’ in many T3SS effectors. ^17^ We further extended the observation by finding both the exchanging of signal sequences and the fusion of new signal termini to form new effectors with varied length (Figure 4). For the first time as we know, we classified the N-terminal signal sequences of YopJ/P proteins into 4 major profiles with several minor profiles, followed by recursively searching proteins with the similar N-terminal profiles. The splitting and re-fusion of effector N-termini and the fusion of new N-termini to other effectors, indicated that the N-termini of T3SS effectors could be considered as independent signal domains, which could be further categorized into different T3SS signal families. This would greatly improve the identification of new T3SS effectors with higher confidence, as there is always a challenge to identify signals with wide sequence divergence. The N-termini of *Bartonella* YopJ/P proteins were not used for HMM profile training, because no T3SS was detected from any *Bartonella* strain and the selection pressure was possibly reduced so that the N-terminal sequences were not recognizable by a T3SS.

A comprehensive list was generated of YopJ/P effectors, effector domain containing proteins and proteins with YopJ/P N-terminal profiles. However, most of these proteins have not been confirmed experimentally. Meanwhile, possible function of the proteins other than T3SS effectors should be investigated especially for those intra-species conserved ones in *Bartonella* or other species. The proposed method is also applicable to other effector families. Hopefully, we can curate the full list of family members of currently verified effectors, effector domains and signal sequences. It will provide us with a large number of new effectors, and help us better understand the pathogenic or symbiotic potential of the bacteria.

## Materials and Methods

### 1 Datasets

Verified YopJ/P effectors were downloaded from T3DB. ^29^ The bacterial genome sequences and gene/protein annotation were downloaded from Genome, GenBank and Protein databases of National Center for Biotechnology Information (http://www.ncbi.nlm.nih.gov/). The full bacterial genome-encoding proteome were put together.

### 2 Hidden Markov Models

The protein sequences were pre-aligned with Clustalw (http://www.ebi.ac.uk/). The alignment results were used for Hidden Markov Model (HMM) profile training with HMMer (http://hmmer.janelia.org), which was further used for protein screening with the built profile.

### 3 Recursive sequence alignment

The recursive sequence alignment strategy was shown in Figure 5. It contained three phases. In Phase 1, The full-length protein sequences of initial known effectors were put together for HMM profile training. Two recursive steps were involved. In Step 1, the effector proteins were directly aligned against all bacterial genome-encoding proteomes with blastp (http://www.ncbi.nlm.nih.gov), and meanwhile the HMM profile was searched from the bacterial proteomes. In Step 2, the representative homologs obtained in each genus were aligned for a second time against the complete bacterial proteome with blastp. Similarly, the homologs obtained through the first step were used for HMM profile updating, and the new profile was screened again in the bacterial proteomes in the second step. All the homologs were collected, curated and redundancy filtering, forming the set of full-length homologs.

**Figure 5.**
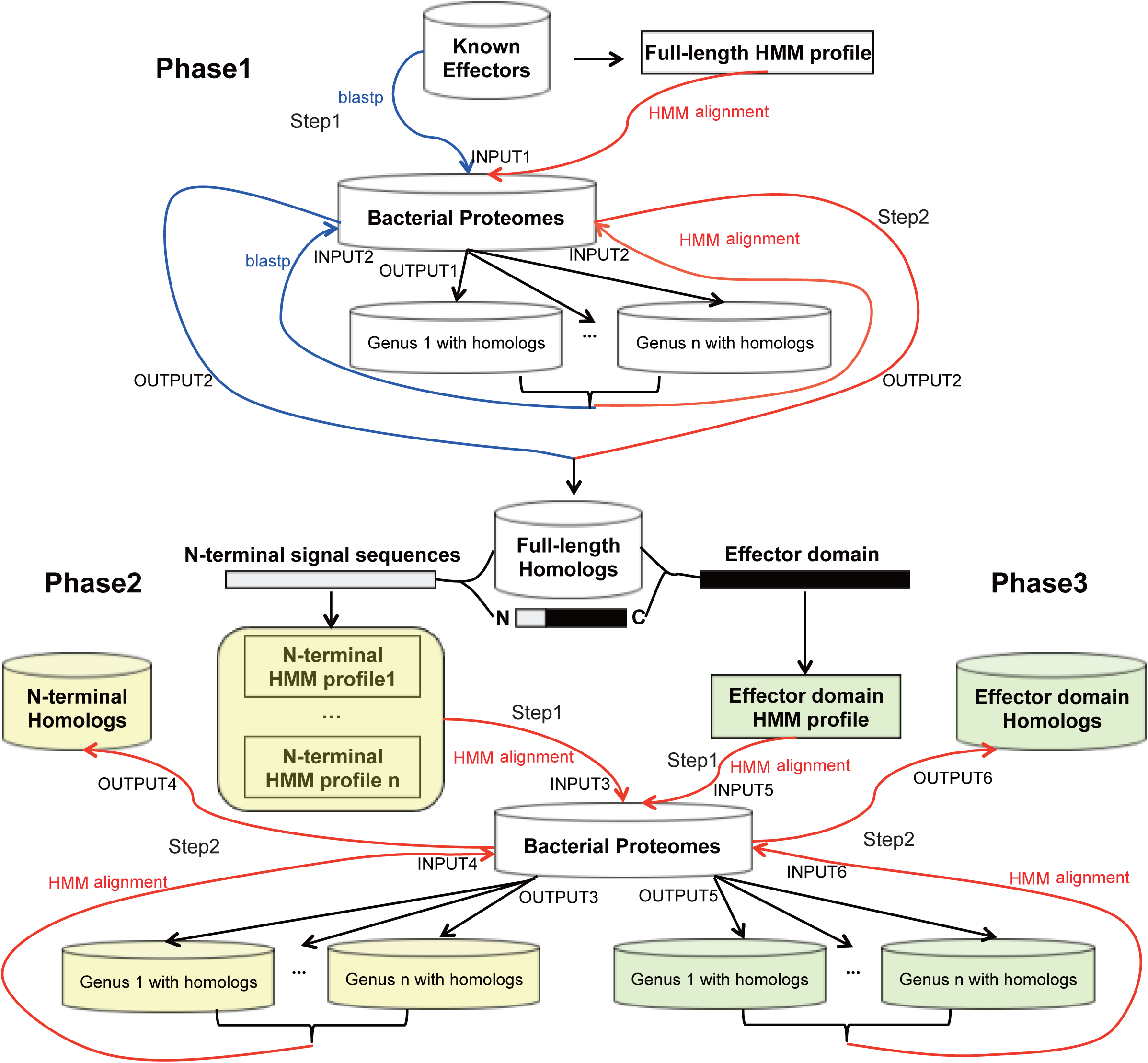
Identification of T3SS effectors by a two-step recursive HMM-based sequence alignment strategy. The identification procedure was separated into three phases: identification of full-length homogs, effector domain homologs and N-terminal signal homologs. For each phase, two successive steps were performed. In the first step, the initial limited number of effectors were used for HMM profile building and homolog searching, and in the second step the HMM profiles were refined with the results of the first step and the homologs were searched again. For each phase, the pipeline was according to the arrowed lines and the order of ‘INPUT1 -> OUTPUT1 -> INPUT2 -> OUTPUT2’ (Phase 1), ‘INPUT3 -> OUTPUT3 -> INPUT4 -> OUTPUT4’ (Phase 2) and ‘INPUT5 -> OUTPUT5 -> INPUT6 -> OUTPUT6’ (Phase 3) respectively as shown in the figure. Blastp and HMM alignment were show in blue and red, respectively. Refer to “Materials and Methods” for details.

Each full-length YopJ/P homolog was split into the N-terminal signal sequence (first 100 amino acids) and the remaining protein sequence containing the C-terminal effector domain. In Phase 2, the N-terminal signal sequences were pre-aligned and classified based on the multiple sequence alignment results. For each cluster, the length was refined, and an HMM profile was built. The profiles were aligned against the bacterial proteome. Only the proteins with similar profile in the N-termini were collected. The N-termini of homologs with the same 1^st^-step profile were aligned again, the length and profile was refined, and the updated profile was aligned against the bacterial proteomes. Proteins with similar N-terminal profiles were collected. In Phase 3, effector domains were aligned and used for domain profile training. The profile was further aligned against the bacterial proteome and the homologs with the effector domain were captured. The homologs from different genera were used for HMM profile updating and then the new profile was screened from the proteomes for another time.

### 4 Detection of Type III Secretion System apparatus proteins

The ten core apparatus proteins, SctC/D/J/N/Q/R/S/T/U/V, were downloaded from T3DB database. ^29^ For each protein family, a HMM profile was built. Each HMM profile was aligned against the target genome-encoding proteomes. Genome coordinates of the genes encoding the apparatus proteins were recorded, and the genomic adjacency was delineated and compared with other known T3SS genes.

## Acknowledgments

The research was supported by a National Natural Science Foundation of China fund to YW (NSFC 81301390) and a National High Technology Research and Development Program (‘863’ Program) to ZG (SS2014AA022210-04).

---

**Supplementary File 1. YopJ|P homologs and proteins with YopJ|P effector domains.txt**

**Supplementary File 2. New effectors with YopJ|P signal sequence profiles.txt**

